# Sustained Effects of Developmental Exposure to Inorganic Arsenic on Hepatic Gene Expression and Mating Success in Zebrafish

**DOI:** 10.1101/2023.07.27.550857

**Authors:** Abigail Ama Koomson, Patrice Delaney, Kirsten C. Sadler

**Affiliations:** Program in Biology, New York University Abu Dhabi, Saadiyat Island, United Arab Emirates

**Keywords:** Zebrafish, Arsenic, Liver, Developmental Origins of Health and Disease

## Abstract

The impacts of exposure to the pervasive environmental toxicant, inorganic arsenic (iAs), on human and fish health are well characterized and several lines of evidence suggest that some impacts can manifest years after exposure cessation. Using a developmental exposure protocol whereby zebrafish embryos were exposed to 0.5 and 1.5 mM iAs from 4-120 hours post fertilization (hpf) and then was removed, we investigated the sustained effects of iAs on gene expression in the liver, survival, reproductive success, and susceptibility to iAs toxicity in the subsequent generation. Developmental exposure to iAs has massive effects on the hepatic transcriptome, with 23% of genes differentially expressed at the end of exposure at 120 hpf, and some of these genes remain deregulated in the liver 9 months after iAs was removed. Developmental exposure to 1.5 mM iAs, but not 0.5 mM, increased mortality through 3 months by over 50%. Adults that were developmentally exposed to 0.5 mM iAs had reduced mating success, but their offspring had no differences in observable aspects of development or their susceptibility to iAs toxicity. This demonstrates that developmental exposure of zebrafish to iAs reduces long-term survival, reproductive success and causes sustained changes to gene expression in the liver.

**SUMMARY STATEMENT:** This work investigates the long-term effects of developmental exposure to inorganic arsenic (iAs) using zebrafish. Months after iAs-exposure was terminated, we found increased mortality, changes in gene expression in the liver and decreased mating success.

## INTRODUCTION

Exposure to inorganic arsenic (iAs) causes a wide range of adverse health outcomes in humans as well as in fish and wildlife (Ahmed et al., 2013; Amuno et al., 2020; Bears et al., 2006; Boyle et al., 2008; Han et al., 2019). iAs is a ubiquitous element naturally present in the bedrock that leaches into water used by humans and as a habitat for aquatic life. Fish and other aquatic species are among the most vulnerable to toxicants as they are chronically immersed in contaminated water and the effects on their health and reproductive success has major impacts for food security and the global ecosystem.

Chronic exposure to iAs in humans causes skin lesions, cardiovascular diseases, respiratory disorders, neurodevelopmental issues, cancer and liver disease (Chen et al., 2023; Frediani et al., 2018; Martinez et al., 2011; Moon et al., 2017; Sanchez et al., 2018; Smeester and Fry, 2018; Xie et al., 2023). We and others have shown that several of the effects of iAs on the liver in mammals also occur during acute exposure of zebrafish (Bambino et al., 2018; Delaney et al., 2023; Delaney et al., 2020; Hallauer et al., 2016; Lam et al., 2006; Li et al., 2016; Sarkar et al., 2017). Many of these occur even after the exposure to iAs has terminated (Thomas, 2013; Young et al., 2018). Thus, it is important to understand how persistent as well as transient exposure translates to long-term health impacts for humans and aquatic animals.

A wealth of data from human and animal studies show that iAs is an agent associated with developmental origins of health and disease (Smeester and Fry, 2018; Young et al., 2018). In humans, exposure to iAs *in utero* during childhood increases risk of several diseases (Farzan et al., 2013), including neurological and cognitive defects, developmental delays (Raqib et al., 2009; Rosado et al., 2007; Tolins et al., 2014; Wasserman et al., 2007), and risk of infection (Farzan et al., 2013; Rahman et al., 2011; Raqib et al., 2009). A case study in Chile showed that early life arsenic exposures increased the rate of liver cancer mortality (Liaw et al., 2008), and these effects persisted in adulthood, with individuals more likely to develop lung and bladder cancer even 40 years after iAs was removed from water sources (Steinmaus et al., 2013). Animal studies also demonstrate that developmental exposure to iAs can have long lasting health impacts. For instance, *in utero* exposure to iAs in mice significantly increased liver and other cancers (Waalkes et al., 2006) and zebrafish exposed to iAs produced offspring that were smaller than controls (Hallauer et al., 2016) and had generational effects on motor activity (Valles et al., 2020). Thus, understanding the effects of iAs on human and wildlife health requires the use of a range of animal models for these studies.

Several studies have investigated the acute and chronic effects of iAs in fish, but few have investigated the long-term or generational effects of developmental exposure. Zebrafish are a widely used model for studying environmental toxicants (Aluru, 2017; Bambino and Chu, 2017; Gamse and Gorelick, 2016), offering many advantages due to their external fertilization, accessibility to large sample sizes, outbred backgrounds, rapid development and transparency of embryos and larvae. Similar to mammals, the zebrafish liver is the primary site of iAs toxicity in zebrafish, and exposure of both larvae and adults causes hepatotoxicity (Bambino et al., 2018; Carlson et al., 2013; Carlson and Van Beneden, 2014; Lam et al., 2006; Li et al., 2016; Ramdas Nair et al., 2022; Sarkar et al., 2017; Yang et al., 2019). As in mammals, iAs toxicity is attributed to cell stress responses, including oxidative and proteostatic stress (Delaney et al., 2020; Fuse et al., 2016; Lam et al., 2006; Ramdas Nair et al., 2022; Xu et al., 2013). We optimized the experimental parameters for using zebrafish in toxicology studies (Ramdas Nair et al., 2021) and extended the use of this model for studying iAs toxicity. We discovered that, like humans, iAs accumulates in the zebrafish liver and that the liver is a major source of iAs toxicity (Bambino et al., 2018; Delaney et al., 2023). We identified developmental time points between 4-5 days post fertilization (dpf), which coincides with liver development, as the prime window of susceptibility to iAs exposure (Delaney et al., 2020). Other studies demonstrated that exposure to iAs during critical developmental stages negatively impact survival, growth, behavior, and has heritable, detrimental effects (Abu Bakar et al., 2022; Hallauer et al., 2016; Valles et al., 2020). Specifically, developmental exposure to iAs (5 – 72 hpf) reduced long-term survival and affected swimming behavior, increased anxiety behaviors and impaired learning in adults (Abu Bakar et al., 2022). Other work has shown that progeny from adults exposed to low doses of iAs for 6-months were small (Hallauer et al., 2016) and that ancestral iAs was associated with reduced motor function in subsequent generations (Valles et al., 2020). While DNA methylation and histone modifications have been proposed to transmit the long-term effects of many toxicants, including iAs (Pilsner et al., 2012; Valles et al., 2020), the epigenetic basis of immediate, latent and heritable consequences of developmental iAs-exposure has not been elucidated in any species.

Here, we use developmental exposure of iAs in zebrafish to investigate the effects on survival, reproductive outcomes, liver gene expression and DNA methylation in exposed animals and on embryonic development and arsenic susceptibility of offspring from adults that were exposed developmentally. Animals that were exposed to high levels of iAs during development had reduced survival, and exposure to lower levels had sustained gene expression changes in the liver and reduced mating success. We found no effects on bulk DNA methylation in exposed animals, and the embryonic development and arsenic toxicity was unchanged in the offspring of adults that were developmentally exposed. This suggests that developmental exposure of iAs in zebrafish embryos has significant health and reproductive effects.

## RESULTS

### Zebrafish liver gene expression recovers from acute (4-5 dpf) iAs exposure

To determine whether acute (4-5 dpf) exposure to iAs causes long-lasting effects on the liver after iAs was removed, we developed a treatment scheme whereby wild-type (WT) zebrafish larvae were exposed to 1.5 mM iAs (the approximate lethal concentration at which 50% of larvae die (LC_50_)) and then washed to remove all residual iAs (**Figure 1A**). Larvae were reared to 24 dpf, encompassing a total time of 19 days post iAs wash out. Livers were dissected from treated and untreated controls at 5, 7, 12, 15, and 24 dpf (0, 2, 7, 10 and 19 days post iAs-washout respectively) and 5-7 livers were pooled for gene expression analysis. A subset of genes reflecting liver function (*bhmt, cetp, cp, dbpa, fabp10a, fgb, fgg, lpl and pgm1*) that we previously reported as downregulated following this treatment protocol (Delaney et al., 2023; Ramdas Nair et al., 2022) were analyzed by quantitative real-time PCR (qRT-PCR) at multiple time points after iAs was removed. Immediately following iAs exposure, these genes were downregulated (**Figure 1B**), reflecting hepatotoxicity. At 2 days post-iAs removal, 6 of these genes increased expression, and by 10 days, expression of all genes returned to baseline levels (**Figure 1B**; **Supplemental Table 1**). This demonstrates that livers in zebrafish recover from acute iAs-toxicity within 10 days of removal.

**Figure 1.**
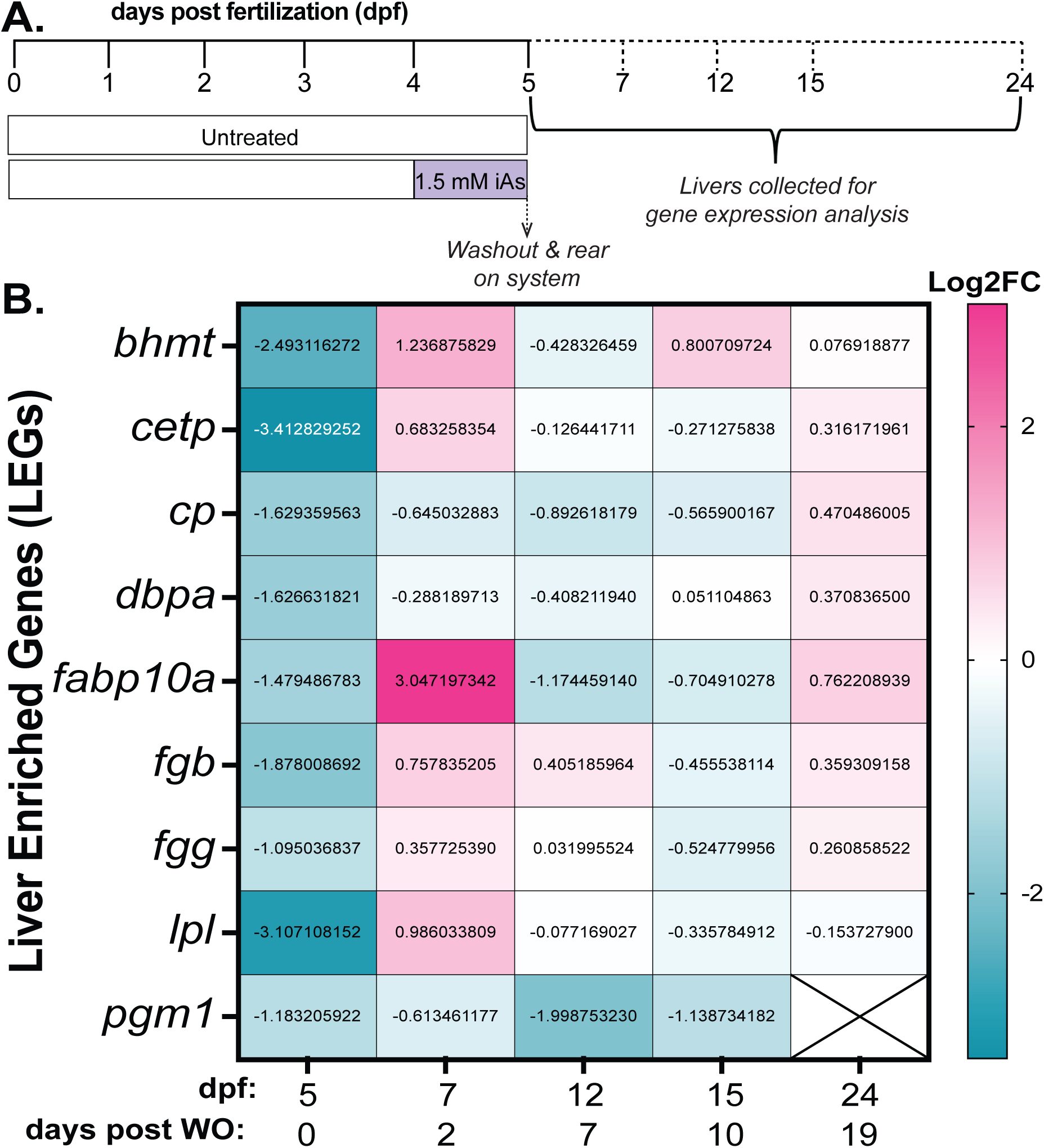
Arsenic responsive genes recover in the liver by 10 days following washout from acute exposure to 1.5 mM iAs. **A.** Treatment scheme of acute treatment (4-5 dpf) to 1.5 mM iAs. At 5 dpf, iAs was washed-out (WO) and zebrafish were reared in the aquaculture system. At the indicated time points, livers were dissected, pooled and analyzed for gene expression. **B.** Heat map of the average log2 fold change (Log2FC) of the expression of genes analyzed in 5-7 livers following iAs WO. The average values from 2 clutches are shown.

### Developmental exposure to iAs induces widespread gene expression changes in the liver

Since we found that there was a relatively rapid recovery of the zebrafish liver to acute iAs exposure, we reasoned that prolonged exposure throughout development could have more potent effects. A previously optimized protocol in which zebrafish were exposed to iAs during major developmental processes including late gastrulation and organogenesis, (4-120 hpf) caused multiple developmental defects, with a lethal concentration 50 (LC_50_) of 1.5 mM (Bambino et al., 2018). Other studies showed that 0.5 mM iAs did not impact the survival or development of zebrafish embryos (Delaney et al., 2020; Li et al., 2009). We therefore used a protocol whereby zebrafish embryos were exposed to 0, 0.5 or 1.5 mM iAs from 4-120 hpf, then it was removed and the animals were reared according to standard aquaculture protocols (**Figure 2A**). As expected, 0.5 mM exposed embryos had no obvious abnormalities while those exposed to 1.5 mM were shorter and had melanocyte clustering over the head (**Figure 2B**). We investigated changes in gene expression in the liver and bulk DNA methylation immediately upon completion of iAs exposure and in subsequent time points, and evaluated survival, morphology, and reproductive success after iAs removal, and then assessed the generational response to iAs toxicity in their offspring (**Figure 2C**).

**Figure 2.**
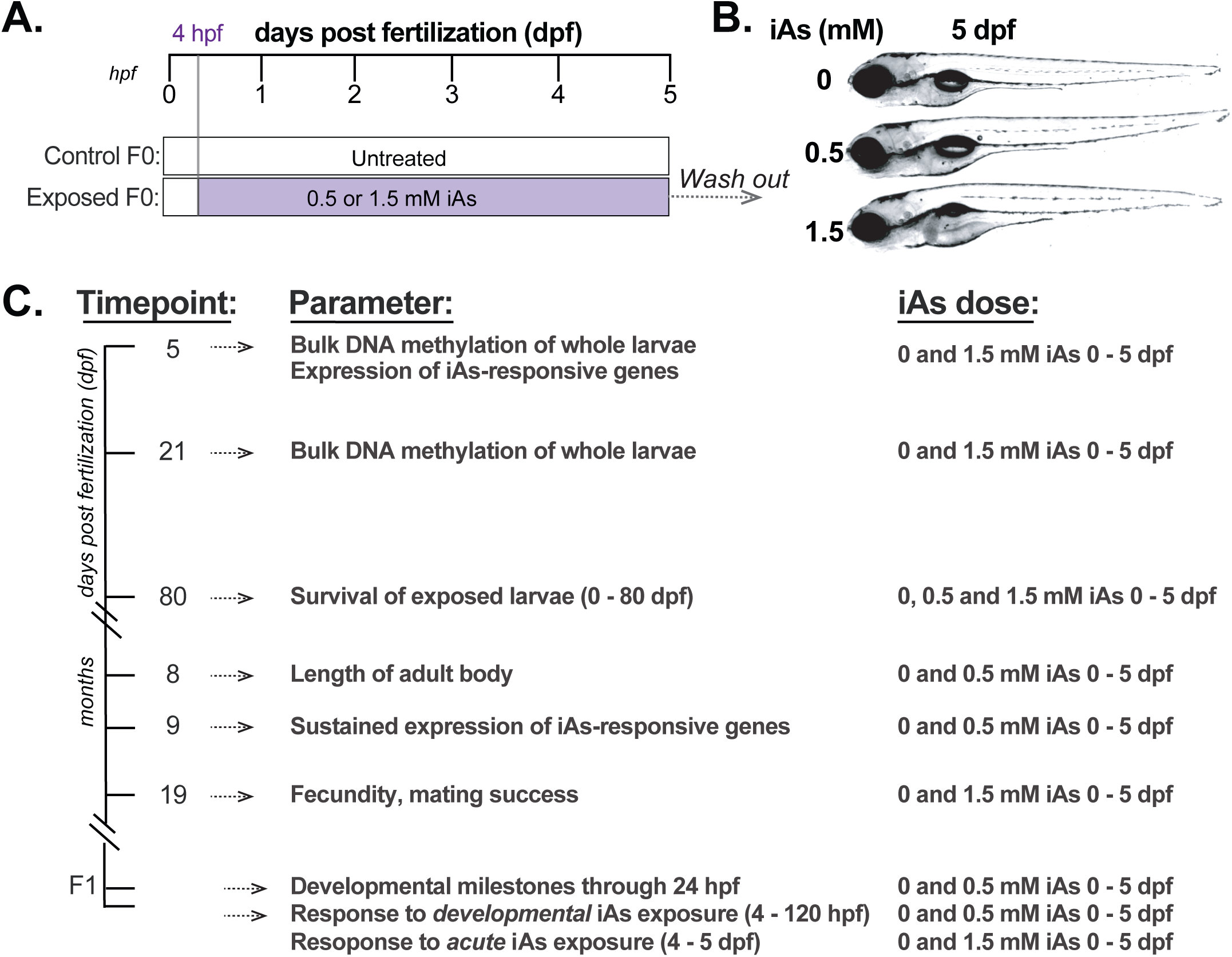
Treatment scheme to evaluate long-term effects of developmental iAs exposure. **A.** Treatment scheme of arsenic exposure where F0 embryos were exposed to 0, 0.5 or 1.5 mM of iAs from 4-120 hpf and then washed out and reared on system. **B.** Representative photos of 120 hpf WT zebrafish exposed to 0, 0.5 or 1.5 mM iAs from 4-120 hpf. **C.** The timeframe of sample collection and parameters assessed.

We analyzed the effects of this exposure protocol on the hepatic transcriptome at 120 hpf, immediately after exposure to 1 mM iAs, using previously published RNAseq data from the liver (Bambino et al., 2018). Nearly 23% of all genes were differentially expressed (padj < 0.05) in these samples **(Figure 3A)**. To determine if developmental exposure to lower (0.5 mM) and higher (1.5 mM) iAs concentrations had the same transcriptomic response, we analyzed a subset of differentially expressed genes (DEGs) that participate in a wide array of hepatic functions, including xenobiotic metabolism, inflammation, and cytokine activity with no two genes overlapping in the same function. We used qRT-PCR to analyze selected upregulated (*aifm4* and *gsto2* (log2FoldChange > 5, padj < 0.05)) and downregulated (*irg1l* and *epob* (log2FoldChange < -3, padj < 0.05)) genes (**Figure 3A**) by quantitative real-time PCR in the liver. We also analyzed the expression of *as3mt* which metabolizes iAs (Ajees and Rosen, 2015), and which we showed is downregulated in the zebrafish liver in response to iAs exposure (Bambino et al., 2018; Delaney et al., 2020). This revealed a similar pattern of expression of these genes in response to 0.5 and 1.5 mM iAs as was found for 1 mM, with a dose-dependent pattern (**Figure 3B**).

**Figure 3.**
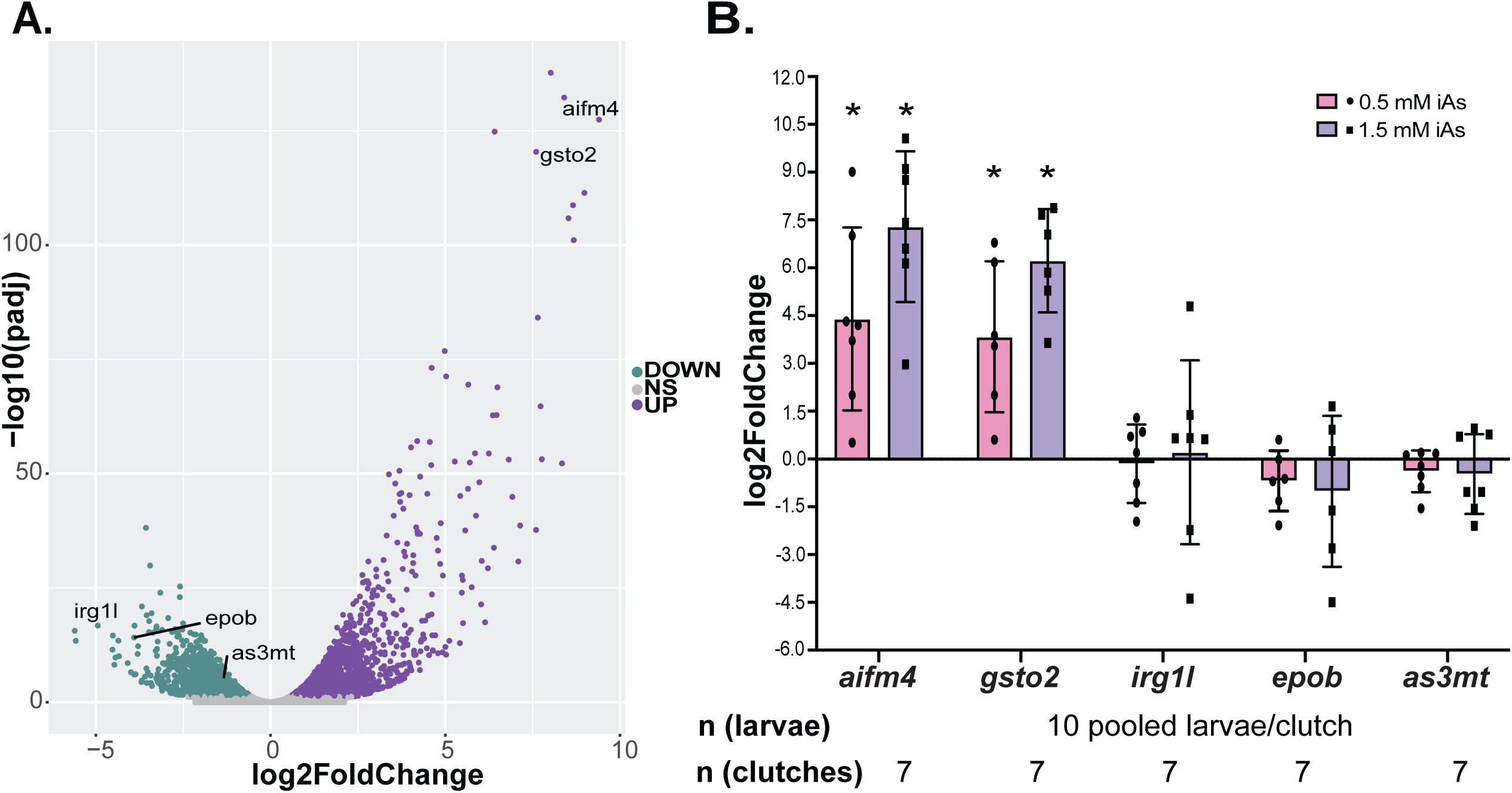
Developmental iAs exposure induces robust gene expression changes in the liver of zebrafish. **A.** Volcano plot of RNAseq data from pools of zebrafish livers following exposure to 1 mM iAs from 4-120 hpf compared to control untreated larvae. The significant (padj<0.05) downregulated genes are labelled in green and upregulated genes are labeled in purple. Selected up and down DEGs for further analysis are labelled. **B.** qPCR analysis from 7 clutches, with 10 livers pooled per clutch collected after 4-120 hpf exposure to 0, 0.5 and 1.5 mM iAs exposure. Values are expressed as the log2 of the fold change for each clutch comparing treated to untreated controls. * denotes p-value < 0.05, t-test.

### Developmental exposure to iAs impacts survival in a dose-dependent pattern

If developmental exposure to iAs has persistent impact on animal health, then we expect to observe increased mortality in a dose-dependent fashion. Tracking the survival of animals that were developmentally exposed to 0.5 and 1.5 mM iAs compared to untreated siblings throughout maturation and adulthood revealed acute mortality immediately following removal of 1.5 mM iAs, and these animals continued to exhibit increased mortality until 40 dpf, with a survival rate of 17% at 50 dpf compared to 55% for controls (**Figure 4A**). In contrast, animals that were developmentally exposed to 0.5 mM iAs had no difference in survival compared to untreated controls (**Figure 4A**) and they showed no changes in overall morphology (**Figure 4B**) or standard length (**Figure 4C**) by 8 months of age.

**Figure 4.**
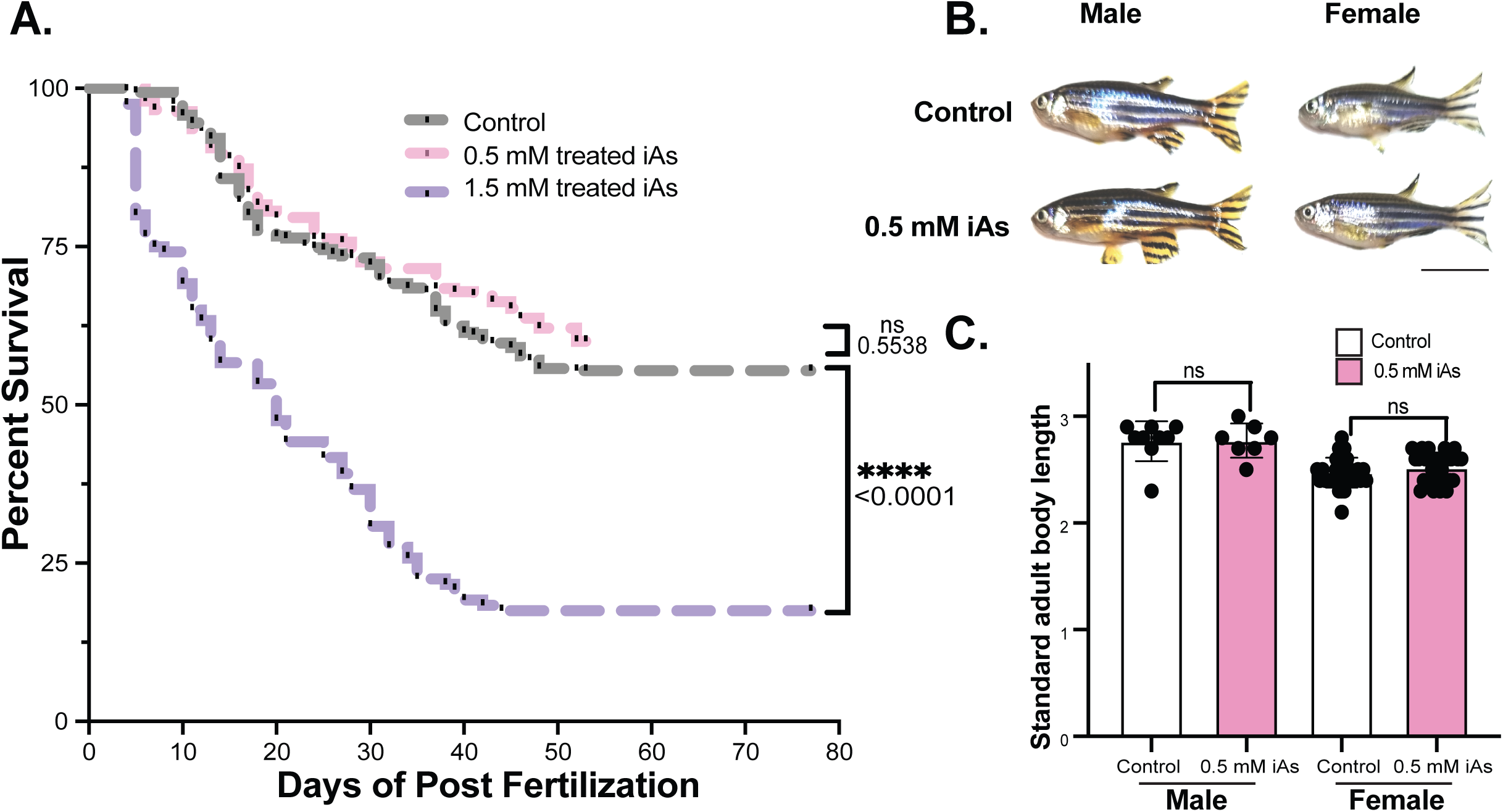
Developmental exposure to iAs negatively impacts long-term survival but not adult morphology. **A.** Survival plot of control (grey), 4-120 hpf 0.5 mM iAs treated larvae (pink) and 4-120 hpf 1.5 mM iAs treated larvae (purple). **B.** Representative images of 8 month old male and female zebrafish that were developmentally exposed to 0.5 mM iAs compared to untreated controls. Scale bar represents 1 cm. **C**. Standard length (cm) of 8 month old males (left) and females (right). Not significant (ns) by Students t-test.

### Developmental exposure to iAs causes persistent gene expression changes in the adult liver

Since developmental exposure to 1.5 mM iAs resulted in high mortality, we reasoned that the animals that survived to adulthood could be adapted to iAs and may not have any long-term effects. We therefore only analyzed adults that were developmentally exposed to 0.5 mM iAs to determine whether developmental exposure to iAs causes sustained gene expression changes in the liver after exposure cessation. Livers collected from 7 males and 7 females (1.5-9 months) which were developmentally exposed to 0.5 mM iAs and an equal number of age matched untreated controls were assessed by qPCR for the same panel of DEGs evaluated at 5 dpf (**Figure 3B**). This showed that the *gsto2* gene which was significantly upregulated immediately following iAs treatment was significantly downregulated in both male and female livers while *aifm4* was significantly down in males and trended down in females, and *epob* trended down in females (**Figure 5A-B**). This demonstrates that developmental iAs-exposure elicits long-term gene expression changes in the liver but these changes are different from the expression changes caused by immediate iAs-exposure.

**Figure 5.**
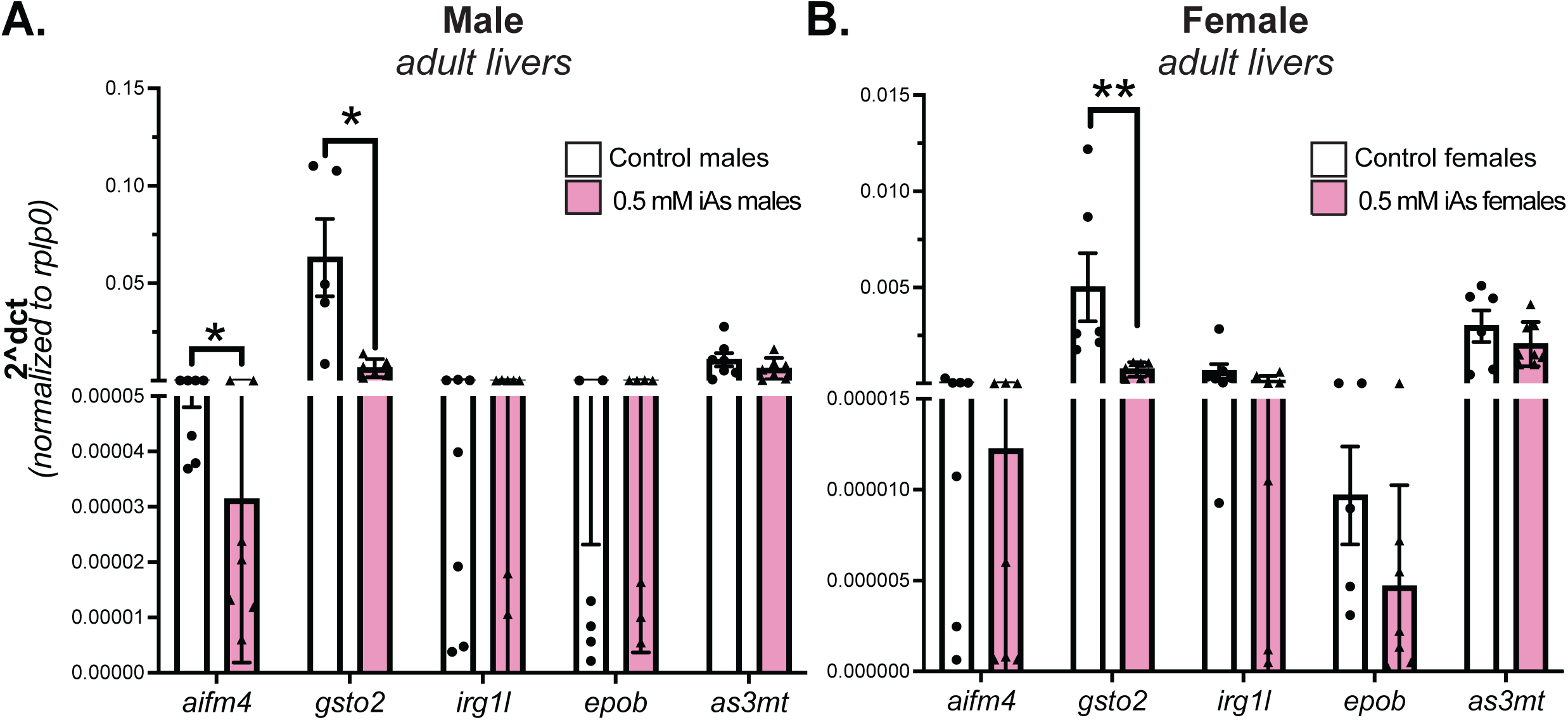
Developmental exposure to iAs sustains differential gene expression in adult livers post wash out. qPCR data from male (**A**) and female (**B**) single livers dissected from age-matched adult zebrafish (1.5 - 9 months old). White bars are from untreated control siblings and pink bars are from the 0.5 mM 4-120 hpf iAs exposed group. n = 7, 2 clutches.

### Developmental Exposure to iAs Does Not Reduce Bulk DNA Methylation Levels

The developmental origins of health and disease theory proposes that epigenetic changes caused by stress or toxicants during development are the mechanism underlying the persistent effects. Some studies indicate that iAs exposure can cause DNA hypomethylation which may contribute in part to long-term iAs-induced disease (Intarasunanont et al., 2012; Nohara et al., 2011). We used biochemical and genetic techniques to examine whether iAs caused DNA hypomethylation during exposure and after cessation. As toxicant exposures generally have very modest effects on DNA methylation levels, we used a high concentration of iAs for these studies to maximize the chance of observing an effect. Whole larvae that were exposed from 4-120 hpf to 0 or 1.5 mM iAs were collected at 120 hpf and then at 21 dpf, 16 days post iAs exposure.

Slot blot analysis of 5-methylcytosine (5-MeC) and double stranded DNA showed that there was no difference in the relative levels of 5-MeC in any sample (**Figure 6A**). We reasoned that since the liver is severely affected by iAs exposure, assessment of 5- MeC whole larvae may not reveal tissue specific changes. We therefore used the transgenic zebrafish reporter of DNA methylation (Goll et al., 2009; Magnani et al., 2021) (*tg(fabp10a:Gal4;cmlc2:EGFP; c269*^o*ff*^*; 10XUAS:dsRed))*, in which EGFP will only be expressed in hepatocytes when the *c269*^o*ff*^ promoter is unmethylated. As a positive control, DNA methyltransferase 1 (*dnmt1)* mutant zebrafish have robust DNA hypomethylation and bright GFP expression in the liver, but no GFP was detected in iAs treated larvae (**Figure 6B**). This demonstrates that iAs does not cause bulk DNA hypomethylation in zebrafish.

**Figure 6.**
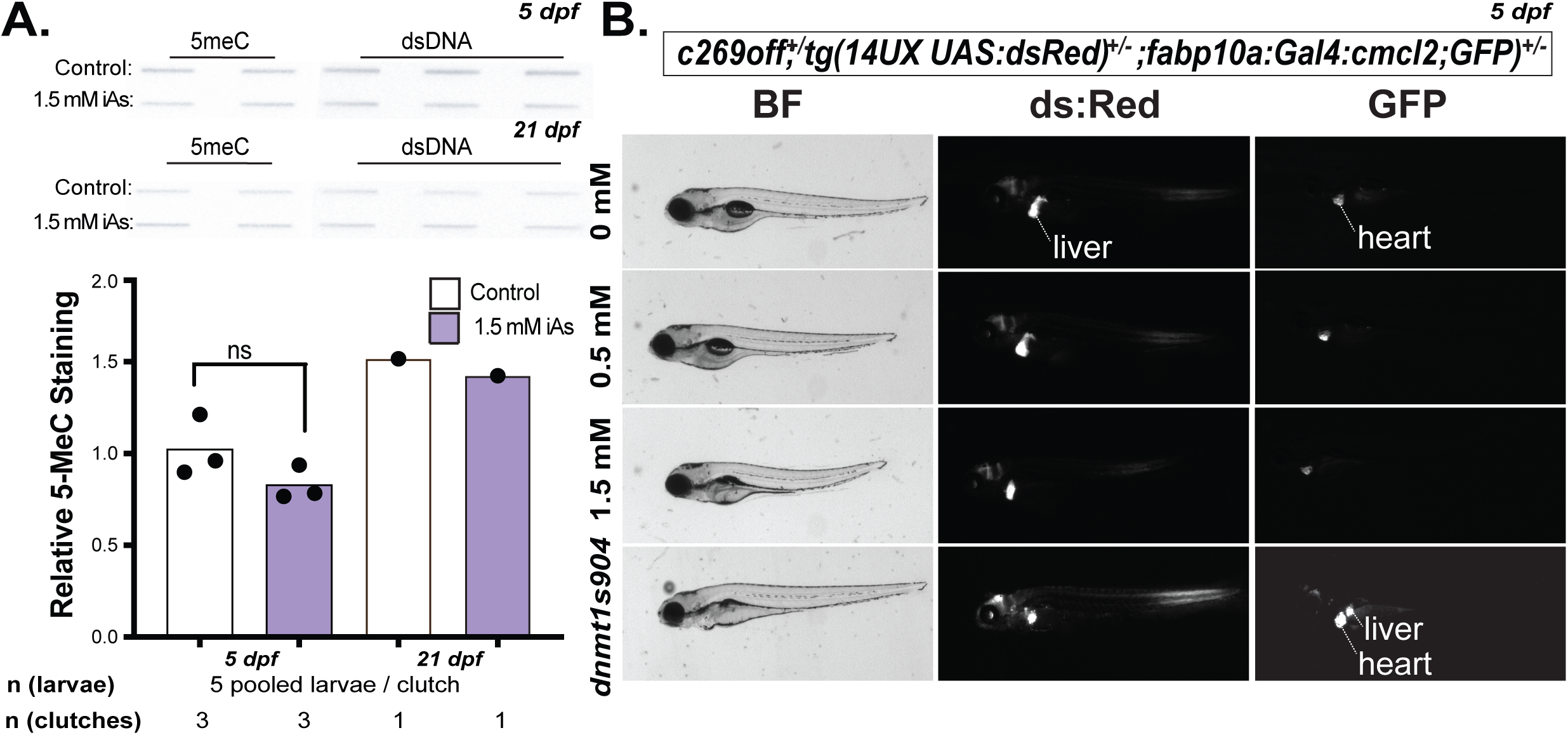
Developmental exposure to iAs does not cause DNA hypomethylation. **A.** Slot Blot hybridization from bulk DNA isolated from whole zebrafish after 0 and 1.5 mM iAs (4-120 hpf) exposure at 5 and 21 dpf (n= 1-3, 1 clutch). ns = not significant by Students t-test. **B.** Representative brightfield and fluorescent images of 5 dpf *tg(fabp10a:Gal4;cmlc2:EGFP; c269*^o*ff*^*; 10XUAS:dsRed)* zebrafish larvae following 0, 0.5 and 1.5 mM iAs exposure from 4-120 hpf. Heart and liver are annotated. *dnmt1s904* zebrafish larvae included as a positive control for hypomethylation.

### Developmental exposure to iAs reduces mating success in adults

To evaluate whether developmental exposure to iAs impacts mating success, the number of spawned eggs and fertilization rate were assessed for 7 pairs of males and females that were reared from developmentally exposed (0.5 mM iAs) or unexposed control embryos. The number of times each pair produced offspring was scored as negative for no embryos and positive for 1 or more embryo produced and each pair was mated up to 5 times. Nearly all pairs of control animals (7/8) mated at least 50% of the time, with 3 pairs producing embryos in all 5 mating trials. The mating success was significantly lower for pairs that were developmentally exposed to 0.5 mM iAs: 2 of the 8 pairs never mated, and of those that did mate, none were successful in all trials. (**Figure 7A**). Of those pairs that did produce offspring, we observed no significant difference in the number of eggs, fertilization rate or developmental progress between embryos generated from control or iAs exposed pairs (**Figure 7B-C).** A similar pattern was observed for offspring generated from adults developmentally exposed to 1.5 mM iAs **(Supplemental Figure S1A-B).**

**Figure 7.**
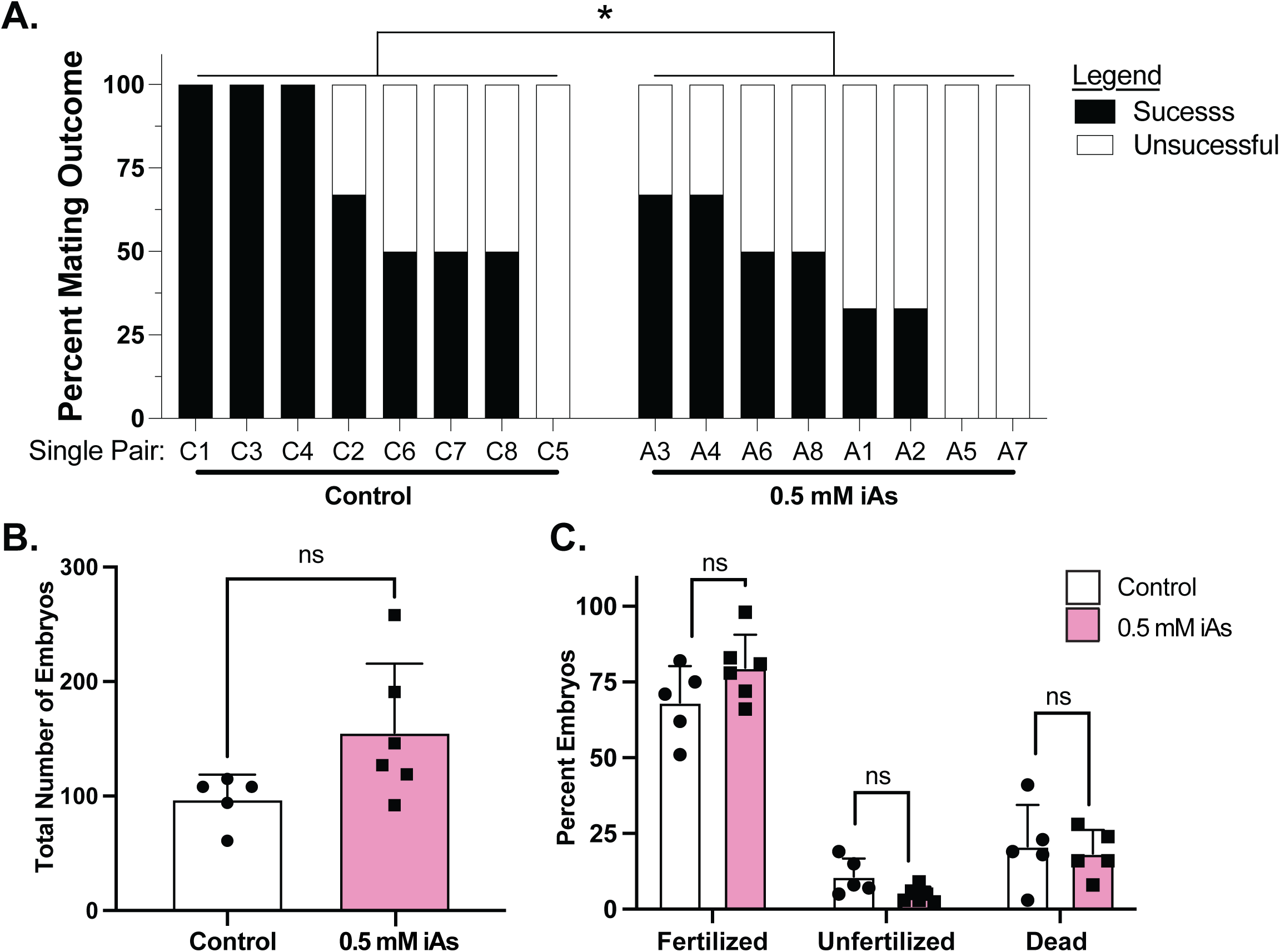
Developmental iAs exposure reduces mating success but does not affect the number of fertilized embryos. **A.** Percent of mating success between 8 single pairs of zebrafish that developed from untreated or 0.5 mM iAs developmentally treated zebrafish. * indicates p<0.05 by Chi Square. **B.** The number of embryos produced from successful matings of control and 0.5 mM iAs developmentally treated zebrafish. **C.** Percent of embryos that were fertilized, unfertilized, and dead from successful mating of control and 0.5 mM iAs developmentally exposed zebrafish.

### iAs toxicity is not changed in offspring from developmentally exposed parents

To determine if parental exposure to iAs during development (F0) alters iAs toxicity in offspring (F1), we exposed F1 embryos generated from parents that were developmentally exposed to 0.5 or 1.5 mM iAs to a range of iAs concentrations from 4- 120 hpf and scored for mortality. There was no difference in the LC_50_ of iAs or the morphology of larvae from control or iAs exposed parents (**Figure 8A-B**, **Supplemental Figure S1C**). Thus, while developmental exposure of zebrafish reduces mating success, the offspring do not have any developmental defects or changes in response to iAs.

**Figure 8.**
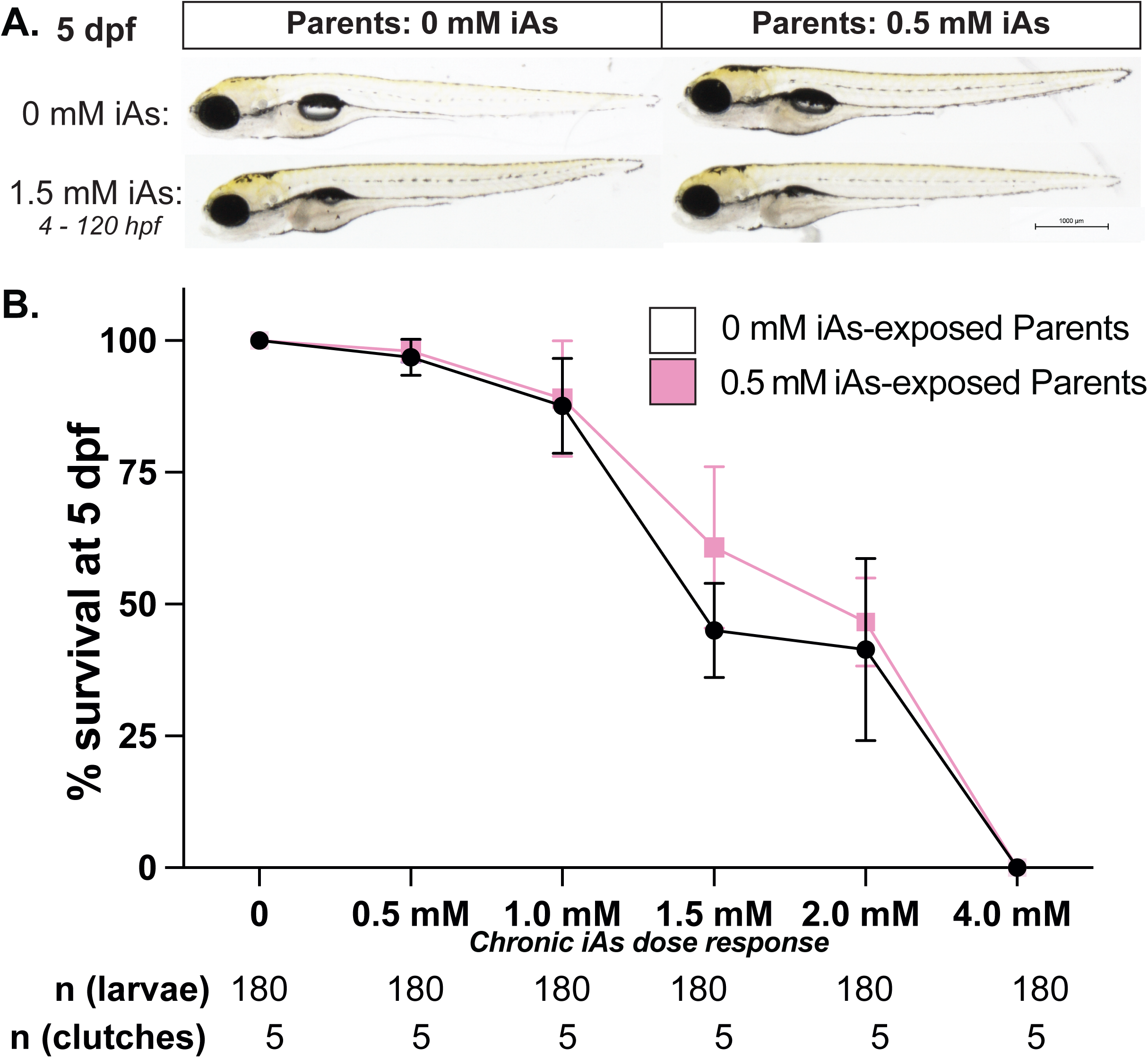
Parental development exposure to iAs does not alter the toxicity of iAs in offspring. **A.** Representative images of 5 dpf F1 larvae produced from incrosses of untreated parents (left) and parents that were developmentally exposed to 0.5 mM iAs (right). F1 were treated to 0 (top) and 1.5 mM iAs from 4-120 hpf (bottom). Bar = 1,000 μM. **B.** Survival curve of F1 produced from incrosses of parents exposed to 0 (black) or 0.5 mM iAs from 0-120 hpf (pink) after 4-120 hpf exposure to 0, 0.5, 1.0, 1.5, 2.0 and 4.0 mM iAs.

## DISCUSSION

The Developmental Origins of Health and Disease field has generated a wealth of evidence that early life toxicant exposure leads to poor health outcomes later in life, and some of these adverse outcomes are passed to offspring (Nicolella and de Assis, 2022). Thus, the effects of toxicants like iAs can cause disease long after the exposure has been terminated. While epigenetic modifications, including DNA methylation, have been proposed as a mechanism of long-term and generational effects of developmental exposures, this has not been established for arsenic. We used zebrafish to investigate how developmental exposure to iAs causes sustained effects on the liver, on reproductive outcomes and generational susceptibility to iAs. We report that the expression of some genes that altered following developmental exposure were also differentially expressed in the adult livers months after iAs was removed. Since we could not detect any changes in DNA methylation levels in the liver following iAs exposure in zebrafish, we speculate that either locus specific DNA methylation changes or other epigenetic modifications underly this long-term effect. Exposure to a high concentration of iAs during development caused acute mortality immediately after removal and increased mortality persisted through adulthood. Lower concentrations of iAs during development which did not affect survival had a significant decrease in mating success but their progeny appeared normal, and had no change in their response to iAs. This data narrows the focus to those aspects of fish biology that are impacted by developmental exposure to iAs to hepatic gene expression changes and reproductive success.

Our finding that iAs induced long-term changes in gene expression in the liver are consistent with other studies demonstrating that developmental exposure to iAs causes changes in gene expression patterns related to neurological disorders. In zebrafish, developmental exposure to iAs (5-72 hpf) caused changes in the expression of genes implicated in autism spectrum disorders at 6 dpf, though only two were significantly upregulated and one was downregulated and it is unknown if this pattern was retained in adulthood (Abu Bakar et al., 2022). Another study showed that the Brain-derived neurotrophic factor *(bdnf)* gene which is implicated in neuronal growth was downregulated in F2 offspring generated from animals that were exposed to iAs from 4 hpf – 150 dpf (Valles et al., 2020). A study in mice showed that pregnant dams fed iAs through drinking water resulted in decreased expression in F2 offspring in some genes, including *BDNF* (Htway et al., 2021). This suggests iAs exposure as a cause of neurological defects in exposed animals and their offspring. Indeed, zebrafish exposed to iAs produced offspring that had behavioral abnormalities (Valles et al., 2020). These changes in behavior provide a possible explanation for the mating success decline in iAs exposed animals as it is possible that they were impaired in their ability to perform or respond to mating rituals.

No studies to date have investigated the persistent effects of iAs on the liver in zebrafish. We identified a subset of genes that are highly iAs responsive in the liver. Interestingly, genes that were highly upregulated after immediate iAs exposure were down regulated in adulthood - *gsto2* and *aifm4*, suggesting a rebound effect on their expression. GSTO2 can conjugate GSH to certain environmental toxins, like iAs, rendering them less toxic and facilitating elimination (Paiva et al., 2010). This is significant as human studies have shown that alleles which decrease GSTO2 function are associated with an increased risk of skin lesions caused by iAs exposure (Luo et al., 2018). This suggests that developmental exposure to iAs could increase susceptibility to iAs toxicity in adulthood. Similarly, *aifm4*, which is predicted to regulate mitochondrial functions, may have an adaptive effect during immediate iAs challenge but become maladaptive over prolonged exposure, potentially explaining the immediate high expression following iAs exposure followed by downregulation in adulthood. Future studies evaluating the entire transcriptome will uncover the complete sustained gene expression response and enable a better understanding of how these contribute to latent iAs-induced diseases.

Our finding that developmental iAs did not reduce bulk DNA methylation immediately after exposure or later after recovery is inconsistent with some studies that suggest DNA hypomethylation as a mechanism of long-term iAs-toxicity (Nohara et al., 2011; Okamura et al., 2019; Reichard et al., 2007). Some studies show that iAs depletes the methyl donor that is used for DNA methylation (Reichard et al., 2007), but we have not evaluated that parameter in our studies. While our data indicate that the transcriptomic changes observed in response to iAs exposure are not attributed to bulk DNA hypomethylation, it is possible that locus specific changes or other epigenetic modifications occur in response to iAs exposure.

Our finding of reduced mating success in adults that were exposed to iAs during development has implications for the fish populations exposed to iAs in their environment. Additionally, this data expands on the studies on iAs exposure and reproductive health outcomes. Developmental iAs exposure in rodents has significant negative effects on sperm quality and quantity (Kim and Kim, 2015; Lima et al., 2018; Pant et al., 2001) and iAs exposure in women has been linked to a wide range of reproductive health issues (Malin Igra et al., 2023; Rahman et al., 2017; Rahman et al., 2021; Tofail et al., 2009). This study provides a model to investigate the mechanism of iAs-induced reproductive outcomes, although it is unclear to what extent the mating success decline in iAs exposed animals can be attributed to defects in gametogenesis or to behavior. Future investigations into whether the reduction in mating success is attributed to male, female or both sexes in zebrafish will help identify groups that are most at risk to long-term impacts in fertility.

## MATERIALS & METHODS

### Zebrafish husbandry and embryo rearing

All procedures were approved by and performed in accordance with the New York University Abu Dhabi Institutional Animal Care and Use Committee (IACUC) guidelines. Adult wild type (WT; ABNYU, TAB5, and BNY14) and transgenic lines *(Tg(fabp10a:Gal4;cmlc2:EGFP; c269*^o*ff*^*; 10XUAS:dsRed))* were maintained on a 14:10 light:dark cycle at 28°C. All experiments were conducted in 6-well plates (Corning, USA) with 20 embryos in 10 mL embryo medium, which is prepared in accordance with the Zebrafish Information Network protocol (Westerfield, 2007). Embryos were collected from group matings or single mating pairs, as indicated, within 2 hours of spawning and were reared at 28°C, according to standard conditions.

### iAs exposure in embryos and adult rearing

Embryos were exposed to sodium meta-arsenite (Sigma-Aldrich, USA; henceforth referred to as iAs) by diluting 0.05 M stock solution to final concentrations ranging from 0.5 mM – 4 mM in embryo medium from either 4-120 hpf or 96-120 hpf, as indicated in the text. A detailed overview of how treatment parameters were optimized are previously described (Ramdas Nair et al., 2021). For all experiments, mortality was scored daily, dead embryos and larvae were removed, and morphological parameters were recorded at 120 hpffr.

After 120 hpf, all treated larvae were washed 3 times in fresh embryo media prior to placing on the NYUAD Fish Facility Aquaculture system (Techniplast) alongside untreated control siblings.

### Image Acquisition

For whole mount imaging of live larvae, embryos were anesthetized with 500 µM tricaine (Ethyl 3-aminobenzoate methanesulfonate; Sigma-Aldrich), mounted in 3% methyl-cellulose on a glass slide and imaged on a Nikon SMZ25 stereomicroscope.

### Gene Expression Analysis

Pools of at least 5 livers were microdissected from 120 hpf zebrafish larvae with transgenic marked livers (*Tg(fabp10a:CAAX-eGFP))*. Larvae were anesthetized in tricaine and immobilized in 3% methyl cellulose and the livers were removed using 30- gauge needles. RNA was extracted from livers using TRIzol (Thermo Fisher, 15596026) and precipitated with isopropanol as described (Vacaru et al., 2014). RNA was reverse transcribed with qScript (QuantaBio, 95048-025).

Gene expression was assessed using RNA-seq or quantitative reverse transcription PCR (qRT-PCR). qRT-PCR was performed using Maxima Sybr Green/ROX qPCR Master Mix Super Mix (Thermo Fisher, K0221). Samples were run in triplicate on QuantStudio 5 (Thermo Fisher). Target gene expression was normalized to *ribosomal protein large P0* (*rplp0*) using the comparative threshold cycle (ΔCt) method. Primers for the genes of interest are listed in **Supplemental Table 2**. Expression in treated animals was compared to untreated controls from the same clutch to determine fold change.

### RNA-seq

The publicly available RNA-seq dataset we generated from transgenic (*tg(fabp10a:nls-mcherry)* zebrafish larvae were untreated (control) or exposed to 1 mM iAs from 4-120 hpf was (GSE104953) was reanalyzed. Differentially expressed genes were identified as those with a padj<0.05. Volcano plots were plotted using R package ggplot2.

### Slot Blot

Slot blot was performed using 0.5 ng of gDNA from pools of five 5 dpf or three 21 dpf zebrafish larvae. gDNA was denatured in 400 mM NaOH/10 mM EDTA and blotted onto nitrocellulose membrane (BioRad) in duplicate for 5mC DNA and triplicate for dsDNA using a slot blot apparatus (BioRad). Equivalent volume of DNAse/RNAse-free water (Invitrogen) was loaded instead of genomic DNA as negative control (data not shown). Membranes were incubated 1 h at 80°C, blocked with 5% bovine serum albumin (BSA) in TBST (37 mM NaCl, 20 mM Tris pH 7.5, 0.1% Tween 20), and incubated overnight at 4°C in either anti-dsDNA (Abcam, 1:5000 in 2% BSA in TBST) or anti-5-methyl-cytosine (5mC – Aviva Biosystem clone 33D3, 1:3000 in 2% BSA in TBST). Membranes were washed in TBST and probed with anti-mouse HRP secondary antibody (Promega; 1:2000 in 5% BSA in TBST) for 1 h at room temperature followed by development in ECL (Thermo Fisher Scientific) or Clarity ECL (BioRad). ChemiDoc (BioRad) was used to detect and quantify the chemiluminescent signal. Gel Analyzer (http://www.gelanalyzer.com) was used to perform quantitative densitometric analysis of the signals and ratio between 5mC and dsDNA was plotted for each sample using GraphPad Prism.

### Statistical analysis, rigor and reproducibility

Experiments were carried out on at least 3 clutches of embryos in most cases and at least 3 adult animals, with all replicates indicated. Reproducibility was assured by carrying out key experiments, including phenotype scoring by independent investigators. Data are presented as normalized values. Statistical tests were used as appropriate to the specific analysis, including Student’s T-test, ANOVA and Chi Square (Fisher’s exact test) using Graphpad Prism Software.

## Acknowledgements

We are grateful for the contributions of Shashi Ranjan for expert fish care and guidance on the project, Sumeet Singh and Nouf Khan for mentorship and critical discussions, and Kathryn Bambino for preliminary data. Gratitude to Bhavani P. Madakashira for proofreading the manuscript.

## Competing interests

No competing interests declared.

## Funding

This research was funded in part by the NYUAD Faculty Research Fund (AD188 to KCS), the NYUAD Division of Science and the Office of Undergraduate Research.

## Diversity and inclusion statement

This work represents a collaborative effort between all authors, with a portion of the senior capstone project of an undergraduate student at NYUAD (AAK) and a doctoral student (PD). All the authors are women.

## SUPPLEMENTAL FIGURE LEGENDS

**Supplemental Figure 1.**
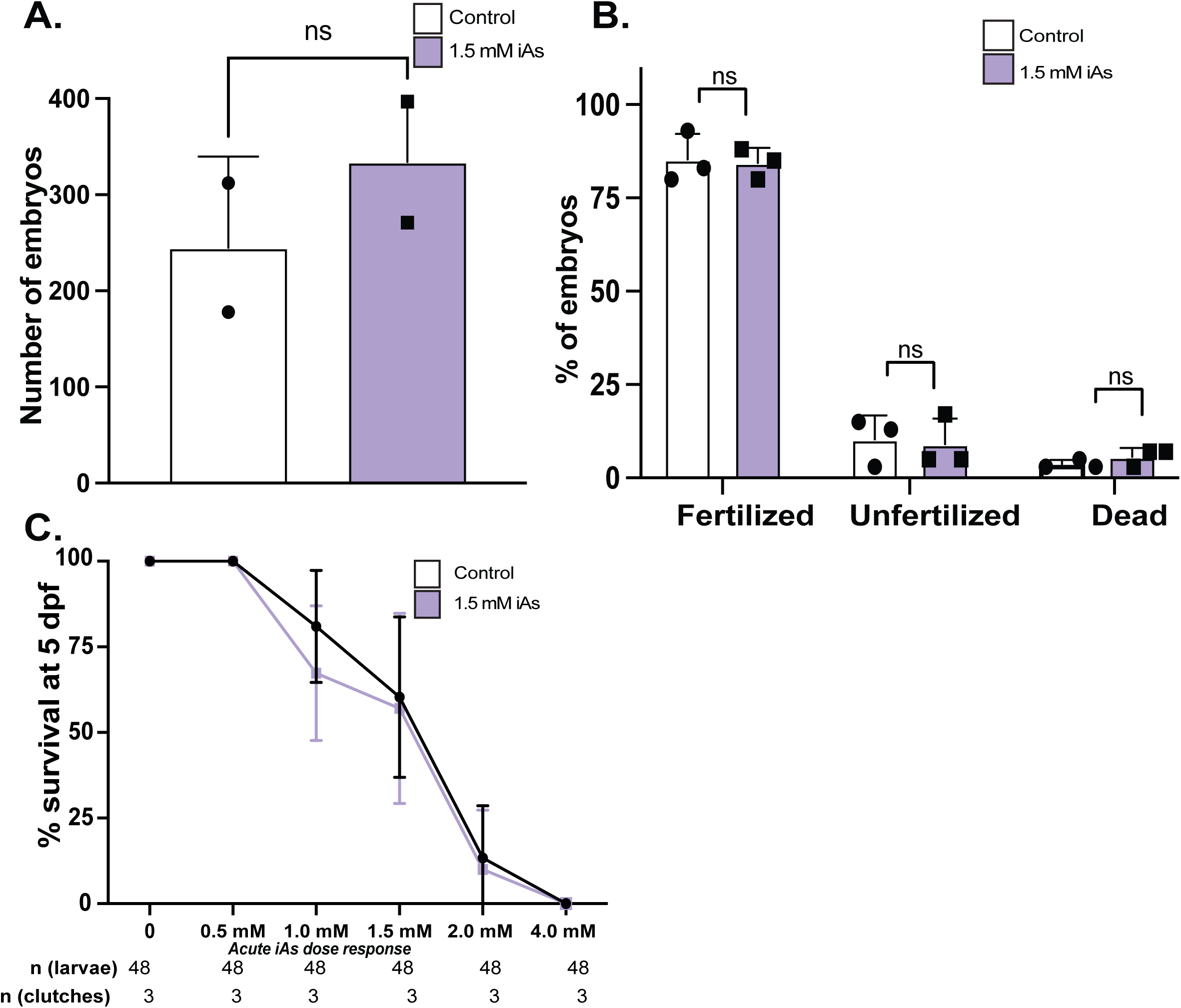
Developmental exposure to 1.5 mM iAs does not alter number of quantity or quality of embryos or F1 sensitivity to iAs. **A.** The number of embryos produced from successful matings of control and 1.5 mM iAs developmentally treated zebrafish. **B.** Percent of embryos that were fertilized, unfertilized, and dead from successful mating of control and 1.5 mM iAs developmentally exposed zebrafish. **C.** Survival curve of F1 produced from incrosses of parents exposed to 0 (black) or 1.5 mM iAs from 0-120 hpf (purple) after 96-120 hpf exposure to 0, 0.5, 1.0, 1.5, 2.0 and 4.0 mM iAs.

## Notes

### Competing Interest Statement

The authors have declared no competing interest.

